# Daytime variation in SARS-CoV-2 infection and cytokine production

**DOI:** 10.1101/2020.09.09.290718

**Authors:** Aïssatou Bailo Diallo, Laetitia Gay, Benjamin Coiffard, Marc Leone, Soraya Mezouar, Jean-Louis Mege

## Abstract

S. Ray and A. Reddy recently anticipated the implication of circadian rhythm in severe acute respiratory syndrome coronavirus 2 (SARS-CoV-2), which is the causative agent of the coronavirus disease (Covid-19). In addition to its key role in the regulation of biological functions, the circadian rhythm has been suggested as a regulator of viral infections. Specifically, the time of day of infection was found critical for illness progression, as has been reported for influenza, respiratory syncytial and parainfluenza type 3 viruses. We analyzed circadian rhythm implication in SARS-CoV-2 virus infection of isolated human monocytes, key actor cells in Covid-19 disease, from healthy subjects. The circadian gene expression of *Bmal1* and *Clock* genes was investigated with q-RTPCR. Monocytes were infected with SARS-CoV-2 virus strain and viral infection was investigated by One-Step qRT-PCR and immunofluorescence. Interleukin (IL)-6, IL-1β and IL-10 levels were also measured in supernatants of infected monocytes. Using Cosinor analysis, we showed that *Bmal1* and *Clock* transcripts exhibited circadian rhythm in monocytes with an acrophase and a bathyphase at Zeitgeber Time (ZT)6 and ZT17. After forty-eight hours, the amount of SARS-CoV-2 virus increased in the monocyte infected at ZT6 compared to ZT17. The high virus amount at ZT6 was associated with significant increased release in IL-6, IL-1β and IL-10 compared to ZT17. Our results suggest that time day of SARS-CoV-2 infection affects viral infection and host immune response. They support consideration of circadian rhythm in SARS-CoV-2 disease progression and we propose circadian rhythm as a novel target for managing viral progression.

**Importance:** The implication of circadian rhythm (CR) in pathogenesis of Severe Acute Respiratory Syndrome Coronavirus 2 (SARS-CoV-2) has been recently anticipated. The time of day of infection is critical for illness progression as reported for influenza, respiratory syncytial and parainfluenza type 3 viruses. In this study, we wondered if SARS-CoV-2 infection and cytokine production by human monocytes, innate immune cells affected by Covid-19, were regulated by CR. Our results suggest that time day of SARS-CoV-2 infection affects viral infection and host immune response. They support consideration of circadian rhythm in SARS-CoV-2 disease progression and we propose circadian rhythm as a novel target for managing viral progression.

## Introductory

The implication of circadian rhythm (CR) in pathogenesis of Severe Acute Respiratory Syndrome Coronavirus 2 (SARS-CoV-2) has been recently anticipated (1, 2). The CR regulates physiological processes in living organisms with a period of 24 hours (3). Rhythmicity depends on central and peripheral oscillators whose activity relies on two main feedback loops managed by a clock genes cascade under the regulation of the main clock gene *Bmal1* (3). The host susceptibility to microorganism is likely under control of biological clocks (4). The time of day of infection is critical for illness progression as reported for influenza, respiratory syncytial and parainfluenza type 3 viruses (5–7). We previously reported that CR is a key actor at the interface between infection susceptibility, clinical presentation and prognosis of infection (4, 8).

There are some evidences that enable to anticipate the role of CR in SARS-CoV-2 infection. The absence of *Bmal1* has an impact on intracellular replication of coronaviruses, especially vesicular trafficking, endoplasmic reticulum and protein biosynthesis (9). Knock-out of *Bmal1* markedly decreases the replication of several viruses such as Dengue or Zika (10). Finally, among key proteins involved in SARS-CoV-2 interaction with the host recently published (11), it has been identified 30% of them being associated with circadian pathway (1). Clearly, the evidences of an implication of CR in SARS-CoV-2 infection of human cells are lacking. In this study, we wondered if SARS-CoV-2 infection and cytokine production by human monocytes, innate immune cells affected by Covid-19, were regulated by CR.

## Results and Discussion

We first wondered if the infection of monocytes, innate immune cells affected by Covid-19 (12), obey to circadian oscillations. Every 3 hours during 24 hours, total RNA was extracted and expression of *Bmal1* and *Clock* genes was investigated in unstimulated monocytes as previously described (8). Expression of investigating genes exhibited CR in monocytes with an acrophase (peak of the rhythm) and a bathyphase (trough of the rhythm) at Zeitgeber (German name for synchronizer) Time (ZT)6 and ZT17 (**Fig.1A**). These two time points represent the beginning of the active and the resting periods in humans (13). To assess the involvement of CR in infection of monocytes with SARS-CoV-2, we incubated monocytes with SARS-CoV-2 during the bathyphase (ZT6) and acrophase (ZT17) during 48 hours. Then, viral RNA was extracted to evaluate Covid-19 virus amount and the phagocytosis index was calculated at ZT6 and ZT17 using immunofluorescence. The amount of SARS-CoV-2 virus was higher in monocytes cultured at ZT6 than at ZT17, suggesting a daytime dependent of viral multiplication (**Fig.1B**). This result was strengthened by high phagocytosis index at ZT6 compared to ZT17 (**Fig.1C**). Our data showed for the first time that entry and multiplication of SARS-CoV-2 in human monocytes varies with the time of day. This finding is reminiscent of what has been previously reported with herpes and influenza virus in murine models of infection (6, 7). It is noteworthy that CRs are different in rodents and humans, thus limitating extrapolations to understand pathogenesis of SARS-CoV-2 infection.

**Figure 1.**
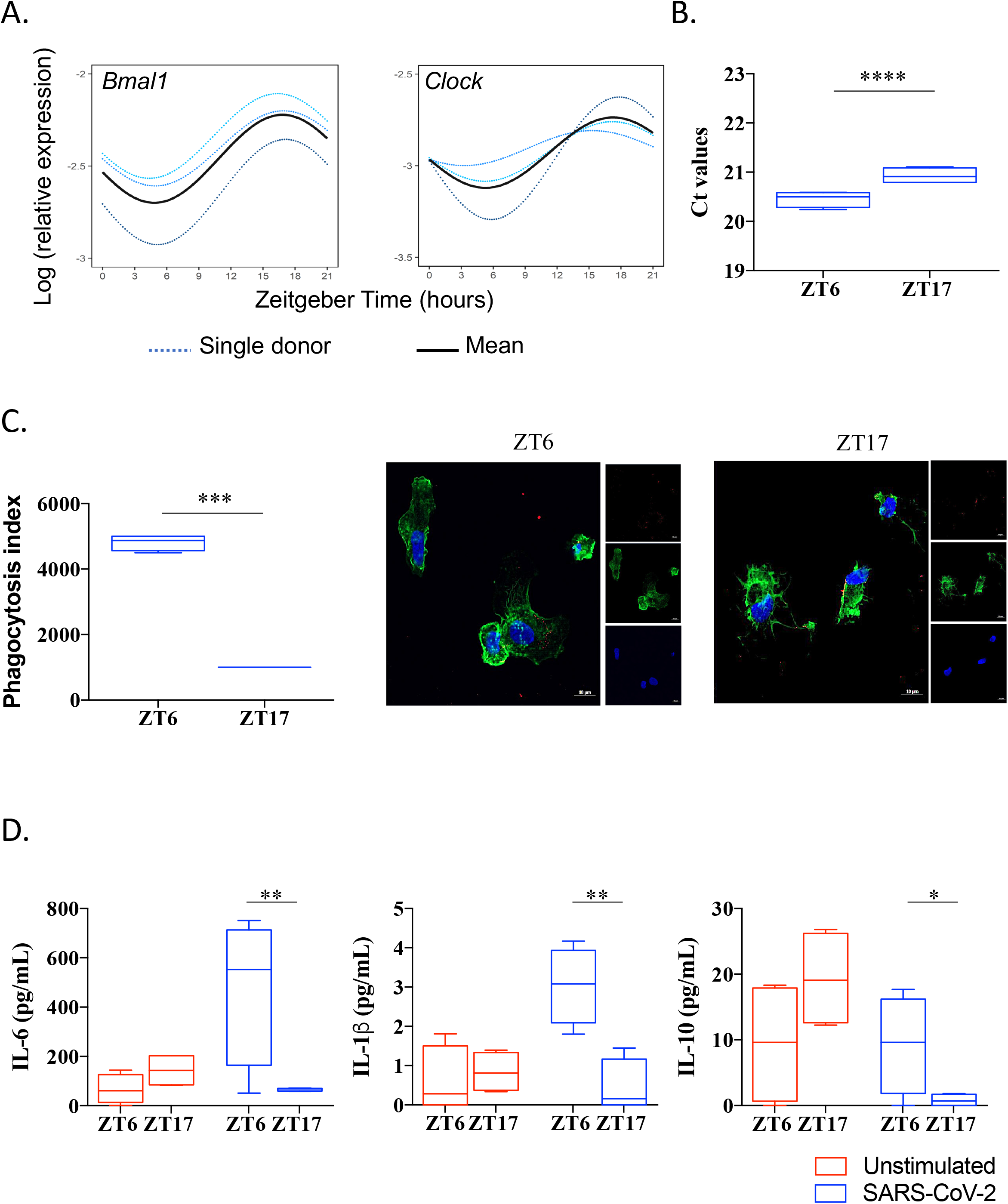
SARS-CoV-2 infection is link to circadian rhythm. (**A**) Circadian rhythm of *BMAL1* and C*LOCK* genes in monocyte using Cosinor model. (**B**) Virus load at ZT6 and ZT17 time. (**C**) Phagocytosis index and representative pictures of monocytes (F-actin in green and nucleus in blue) infected by SARS-CoV-2 virus (red). (**D**) Level of IL-6, IL-1β and IL-10 of unstimulated (red) and infected cells at ZT6 and ZT17.

Covid-19 disease is characterized by runaway immune system leading to a cytokine storm consisting of high circulating levels of cytokines including IL-6, IL-1β and IL-10 (14). We wondered if the interaction of SARS-CoV-2 with monocytes affected cytokine production at two points of the CR. The amounts of IL-1β, IL-6 and IL-10 were significantly increased at ZT6 (**Fig.1D**) when the amount of infection is highest. Hence, the interaction of SARS-CoV-2 with monocytes resulted in distinct cytokine pattern according to daytime.

We demonstrate here that the time day of SARS-CoV-2 infection determines consistently viral infection/replication and host immune response. It is likely that SARS-CoV-2 exploits clock pathway for its own gain. Our findings support consideration of CR in SARS-CoV-2 disease progression and suggest that CR represents a novel target for managing viral progression. This study also highlights the importance of the time of treatment administration to Covid-19 patients since CR was found regulating pharmacokinetics of several drugs (15). Several treatments are proposed to prevent the occurrence of severe forms in Covid-19. They include passive immunization, cytokines, anti-cytokine antibody or corticoids (16). All these candidates affect the immune response known to oscillate during the day and their administration according to CR of SARS-CoV-2. Finally, the well-documented CR disturbance in intensive care units (17) should be considered in the clinical and therapeutic management of patients with severe Covid-19.

## Methods

### Cells and virus

SARS-CoV-2 strain MI6 was cultured in Vero E6 cells (American type culture collection ATCC® CRL-1586™) in Minimum Essential Media (Life Technologies, Carlsbad, CA, USA) supplemented with 4% fetal bovine serum (FBS), as previously described (18).

Human monocytes were isolated from peripheral blood mononuclear cells from healthy donors (convention *n*°*7828*, Etablissement Français du Sang, Marseille, France) following CD14 selection using MACS magnetic beads (Miltenyi Biotec, Bergisch, Germany) as previously described (19). Monocytes were cultured in Roswell Park Memorial Institute medium-1640 (Life Technologies) containing 10% of FBS, 100 U/mL penicillin and 50 µg/mL streptomycin (Life Technologies). Monocytes were infected with 50 µl virus suspension (0.1 multiplicity of infection (MOI)) during bathyphase (ZT6) and acrophase (ZT17) for 48 hours at 37°C in the presence of 5% CO_2_.

### Circadian gene expression

Total RNA was extracted using the RNA Mini Kit (Qiagen) and a quantitative Real-Time PCR was performed according to the manufacturer’s instructions (MMLV Kit, Life Technologies and Smart SYBRGreen kit, Roche Applied Science). Circadian gene expression was investigated using specific primers targeting *Bmal1* and *Clock* genes (8). Results were normalized using the housekeeping endogenous control *actb* gene (β-actin). The results are expressed according to the appropriate formula: gene expression = Log (2^-ΔCt^) relative expression, with Ct (Cycle threshold), ΔCt = Ct target gene - Ct β-actin. The Cosinor analysis based on an extrapolation from measurements of a few points over 24 hours was used to evaluate the CR of the clock genes *Bmal1* and *Clock*.

### Viral RNA extraction and PCR

Viral RNA was extracted from the infected cells using NucleoSpin® Viral RNA Isolation kit (Macherey-Nagel) and Covid-19 virus detection was performed using One-Step qRT-PCR SuperScript™ III Platinum™ (Life Technologies) targeting the gene E, as previously described (20).

### Immunofluorescence

Infected cells (5. 10^5^ cells/well) were fixed and incubated with phalloidin-555 and 4’,6-diamidino-2-phenylindole (DAPI) to labelled F-actin and nucleus respectively. SARS-CoV-2 virus was labelled using first an anti-SARS-CoV-2 antibody (Spike protein, Thermo Fischer) and then a secondary anti-rabbit Alexa 647 (Thermo Fisher).

Pictures were acquired using confocal microscopy (LSM 8000 Airyscan confocal microscope, x 63, oil objective) and the phagocytosis index was calculated according to the following formula: percentage of phagocytosis ((average number of infected cells x 100)/total number of counted cells) x average number of particles or viruses/cells).

### Immunoassays

Levels of interleukin (IL)-6, IL-1β and IL-10 were measured in cell supernatants using an enzyme-linked immunosorbent assay technique (R&D systems). The sensitivity of the assays was (pg/ml) 15.4 for IL-6, 0.125 for IL-1β and 3.9 for IL-10.

### Statistical analysis

Statistical analyses were performed with GraphPad Prism (7.0, La Jolla, CA) and R studio v3.4.0. Continuous variables were expressed as medians ± interquartile, and comparisons between two groups were made using the Mann-Whitney non-parametric test for unmatched data and the Student t-test for matched data. Statistical significance was defined as *P*≤*0*.*05*.

## Author contributions

A.B.D and L.G performed experiments. A.B.D, B.C and S.M analyzed the data. A.B.D, M.L, S.M and J.L.M supervised the work. A.B.D, S.M and J.L.M wrote the manuscript. All authors reviewed and approved the submitted manuscript. All authors reviewed the draft of the manuscript and provided intellectual input.

